# Generation of a mutator parasite to drive resistome discovery in *Plasmodium falciparum*

**DOI:** 10.1101/2022.08.23.504974

**Authors:** Krittikorn Kümpornsin, Theerarat Kochakarn, Tomas Yeo, Madeline R Luth, Richard D Pearson, Johanna Hoshizaki, Kyra A Schindler, Sachel Mok, Heekuk Park, Anne-Catrin Uhlemann, Sonia Moliner Cubel, Virginia Franco, Maria G Gomez-Lorenzo, Francisco Javier Gamo, Elizabeth A Winzeler, David A Fidock, Thanat Chookajorn, Marcus CS Lee

**Affiliations:** Wellcome Sanger Institute, Wellcome Genome Campus, Hinxton, United Kingdom; The Laboratory for Molecular Infection Medicine Sweden and Department of Molecular Biology, Umeå University, Umeå, Sweden; Department of Microbiology and Immunology, Columbia University Irving Medical Center, New York, New York, United States; Department of Pediatrics, School of Medicine, University of California, San Diego, La Jolla, California, United States; Center for Malaria Therapeutics and Antimicrobial Resistance, Division of Infectious Diseases, Department of Medicine, Columbia University Irving Medical Center, New York, New York, United States; Global Health Medicines R&D, GlaxoSmithKline, Tres Cantos, Madrid, Spain; Genomics and Evolutionary Medicine Unit, Centre of Excellence in Malaria Research, Faculty of Tropical Medicine, Mahidol University, Bangkok, Thailand

**Keywords:** *P. falciparum*, DNA polymerase δ mutant, Mutation rate, Mutator, Genetic repertoire, Drug resistance evolution

## Abstract

*In vitro* evolution of drug resistance is a powerful approach for identifying antimalarial targets, however key obstacles to eliciting resistance are the parasite inoculum size and mutation rate. Here we sought to increase parasite genetic diversity to potentiate resistance selections by editing catalytic residues of *Plasmodium falciparum* DNA polymerase δ. Mutation accumulation assays revealed a ∼5-8 fold elevation in the mutation rate, with an increase of 13-28 fold in drug-pressured lines. When challenged with KAE609, high-level resistance was obtained more rapidly and at lower inoculum than wild-type parasites. Selections were also successful with an “irresistible” compound, MMV665794 that failed to yield resistance with other strains. Mutations in a previously uncharacterized gene, PF3D7_1359900, which we term quinoxaline resistance protein (QRP1), were validated as causal for resistance to MMV665794 and an analog, MMV007224. The increased genetic repertoire available to this *“*mutator” parasite can be leveraged to drive *P. falciparum* resistome discovery.

## INTRODUCTION

Antimalarial drug discovery has been actively searching for new or improved medicines to treat and ultimately eliminate malaria. Current front-line artemisinin-based combination therapies (ACTs) for *Plasmodium falciparum* have been compromised by the emergence of less susceptible parasites to both artemisinin and partner drugs in Southeast Asia, an epicenter of antimalarial resistance ^1, 2^. Furthermore, artemisinin resistance is a public health threat to people living in endemic regions worldwide, as exemplified by recent reports of the emergence of Kelch13 mutations in Rwandan and Ugandan isolates that cause reduced artemisinin susceptibility ^3, 4^. Many promising antimalarial compounds with good potency and multi-stage activity have been uncovered using phenotypic-based screening ^5^. However, this approach presents difficulties for lead optimization because of the lack of knowledge of the molecular target. A deeper understanding of the drug target, mode of action and resistance mechanism could lead to the design of better medicines that can withstand drug resistance ^6, 7, 8^. In addition, the drug target can be employed as a molecular marker for genomic epidemiology surveillance in the field to monitor the spread and containment of drug resistance ^9, 10^.

*In vitro* evolution of drug resistance followed by whole-genome analysis has become a key approach for drug target identification by helping define modes of action as well as mechanisms and propensities for resistance ^11, 12, 13^. A typical *in vitro* resistance selection is performed using a parasite inoculum ranging from 10^5^ to 10^9^ parasites, which are exposed to an antimalarial compound at a concentration capable of killing all the parasites to sub-microscopic level ^8, 14, 15^. Recrudescent parasites can then be subjected to whole-genome sequencing to identify the underlying gene responsible for the resistance phenotype. The main obstacle to success is the prolonged or in some cases complete inability to select for resistant parasites, regardless of the selection regime or strain background. This labour- and time-intensive process may thus fail to identify a molecular target or a defined mechanism of action for query compounds. For example, in a set of *in vitro* resistance selections with 48 compounds by the Malaria Drug Accelerator Consortium (MalDA), 23 compounds yielded resistant parasites with resistance observed after 15-300 days ^16^. Compounds that fail to yield resistant parasites after multiple attempts have been termed “irresistible” ^17^. Although there may be multiple possible reasons for compounds to prove “irresistible”, their low propensity for resistance is an attractive quality and thus insights into their mechanism of action would be valuable.

The ability to select for a parasite with a protective mutation depends, at least in part, on an inoculum size with sufficient genetic variation. However, for reasons of technical practicality the maximum inoculum for *in vitro* resistance selections is typically capped at ∼5×10^9^ parasites per flask (∼10% parasitaemia with 3% haematocrit in a 170 mL culture), orders of magnitude less than can occur in an infected human. Larger culture sizes and extended selection times also consume more compound, which may be limiting. Several laboratory strains, as well as field isolates collected from the drug resistance epicenter of western Cambodia, have been shown to have a similar range of mutation rates of around 10^−9^ to 10^−10^ base substitutions per site per asexual life cycle ^18, 19, 20, 21^.

To increase the genetic diversity represented in a given culture volume and potentially shorten the experimental time scale of selection, we used CRISPR-Cas9 to generate a *P. falciparum* mutant Dd2 parasite that had deficient proof-reading activity of the DNA polymerase δ catalytic subunit. We show that this engineered line has an increased mutation rate, lowering the inoculum and shortening the time required to select resistance to KAE609, a compound with a known target ^22^. When challenged with a previously irresistible compound MMV665794 that had failed in selections with wild-type 3D7 and Dd2 parasites ^16, 23^, we were able to obtain multiple resistant clones with mutations in a gene of unknown function, PF3D7_1359900. CRISPR-Cas9 editing of these candidate mutations into wild-type parasites conferred a similar level of resistance to the selected line, demonstrating the role of this gene in resistance to quinoxaline-based compounds. Our results support the potential of this “mutator” parasite to identify new antimalarial targets and understand drug resistance mechanisms.

## RESULTS

### CRISPR editing of DNA polymerase δ

To increase the genetic repertoire of *P. falciparum* parasites in culture, we hypothesized that parasites with impaired 3ʹ-5ʹ proof-reading activity from the catalytic subunit of DNA polymerase δ (PF3D7_1017000) could increase the level of basal spontaneous mutations, based on prior work in yeast and the rodent malaria parasite *Plasmodium berghei* ^24, 25, 26^. The high-fidelity replicative DNA polymerase δ is a major enzyme for lagging-strand synthesis and contains 3ʹ-5ʹ exonuclease activity that can excise misincorporated nucleotides during DNA replication ^27, 28^. The disruption of two conserved catalytic residues of the exonuclease domain of DNA polymerase δ (**Supplementary Figure 1**) leads to impaired 3ʹ-5ʹ proof-reading activity, resulting in reduced fidelity in DNA replication. This causes an increase in nucleotide sequence variation and higher mutation rate in the genome ^29, 30^. The two conserved catalytic residues of the *P. falciparum* 3ʹ-5ʹ proof-reading subunit, D308 and E310, were replaced with alanine using CRISPR-Cas9 in the Dd2 strain background (**Figure 1A and 1B**).

**Figure 1.**
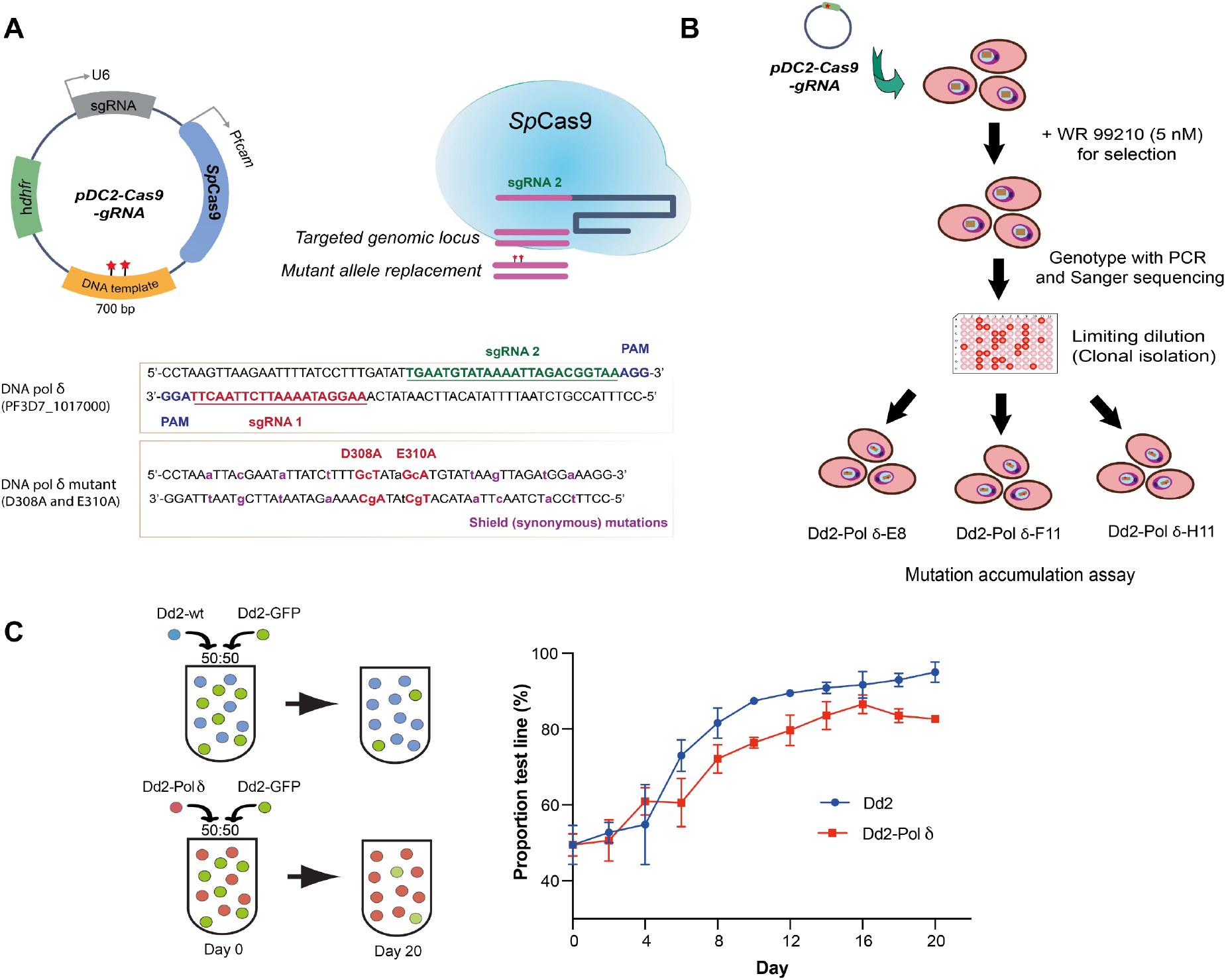
CRISPR-Cas9 editing of the DNA polymerase *δ* proof-reading subunit. **A**) The D308 and E310 residues were replaced by alanine in *P. falciparum* Dd2 by using the pDC2-Cas9-gRNA plasmid containing the sgRNA, Cas9 and donor template. The two sgRNA binding sites and the silent shield mutations are indicated. **B**) CRISPR-Cas9 edited parasites were selected with 5 nM WR99210 and edited clones were isolated by limiting dilution. Three clonal DNA polymerase *δ* mutant parasites (Dd2-Pol*δ*) were selected for whole-genome sequencing and the mutation accumulation assay. **C**) Fitness of the Dd2-Pol*δ* mutant parasite. Competitive fitness assays mixed the fluorescent reference line Dd2-GFP in a 1:1 ratio with either Dd2-WT or Dd2-Pol*δ* clone H11. Growth was determined by flow cytometry every two days for a total of 20 days, with the proportion of GFP-positive parasites compared with total infected RBCs detected by MitoTracker Deep Red. Two independent experiments with technical triplicates were performed, error bars show standard deviation (SD).

*P. falciparum* DNA polymerase δ is predicted to be essential for parasite survival ^31^. To examine whether ablation of the proof-reading function of DNA polymerase δ incurred a fitness cost to the parasite, we performed a competitive fitness assay. Dd2-GFP, an engineered parasite that strongly expresses green fluorescence protein (GFP), was used as a growth reference ^32^. The reference Dd2-GFP line was mixed in a 1:1 ratio with either Dd2 wild-type (Dd2-WT) or the Dd2 DNA polymerase δ mutant (Dd2-Polδ), and the relative proportions of the two lines was measured by flow cytometry every two days for 20 days (∼10 generations). Dd2-Polδ showed only a slightly reduced fitness compared with Dd2-WT based on how quickly each line was able to outcompete the more slowly proliferating Dd2-GFP reference (**Figure 1C**).

### Impaired proof-reading DNA polymerase δ increases single nucleotide variants

To test for changes in nucleotide sequence diversity and mutation rate, we performed a mutation accumulation assay in combination with whole-genome sequencing (**Figure 2A**), comparing Dd2-Polδ with Dd2-WT. Three clones of Dd2-Polδ (E8, F11, H11) and a clone of Dd2-WT were cultured in complete medium continuously for 100 days (∼50 generations) (**Figure 2A**). Parasites were collected every 20 days and clones were isolated by limiting dilution. A total of twelve clones of Dd2-WT and 37 clones of Dd2-Polδ, corresponding to one to three clones per timepoint, were randomly selected for whole-genome sequencing. The genomes of all parasites were mapped to the Dd2 reference genome (PlasmoDB-44_PfalciparumDd2_Genome). The Dd2 core genome comprising coding and non-coding regions was employed as the reference for single nucleotide variant (SNV) calls. The variant surface antigen gene family (*var*) and subtelomeric region of all chromosomes were excluded from the core genome. The genomic coordinates of Dd2 chromosomes were defined in **Supplementary Table 9**. The *de novo* SNVs for each of the clones were determined by comparison with their parental lines on day 0. The number of *de novo* SNVs in Dd2-WT was on average less than 1 SNV per clone in the coding sequence over the 100-day culture period. In contrast, each of the Dd2-Polδ clones had on average 3 - 6 SNVs per clone in coding regions (exome) (**Figure 2B, Supplementary Figure 2A, and Supplementary Tables 1 and 2**). The difference in the number of SNVs in non-coding regions between Dd2-WT and Dd2-Polδ clones was less pronounced (**Figure 2B**). Nonetheless, each of the Dd2-Polδ clones had a greater number of SNVs of all types, distributed across all 14 chromosomes (**Figure 2C and Supplementary Figure 3**). Comparison of base pair substitutions for transition (Ts) and transversion (Tv) events showed a moderate decrease in the Ts:Tv ratio in Dd2-Polδ and an increase in G:C → A:T transitions of 2-4 fold (**Supplementary Figure 4**).

**Figure 2.**
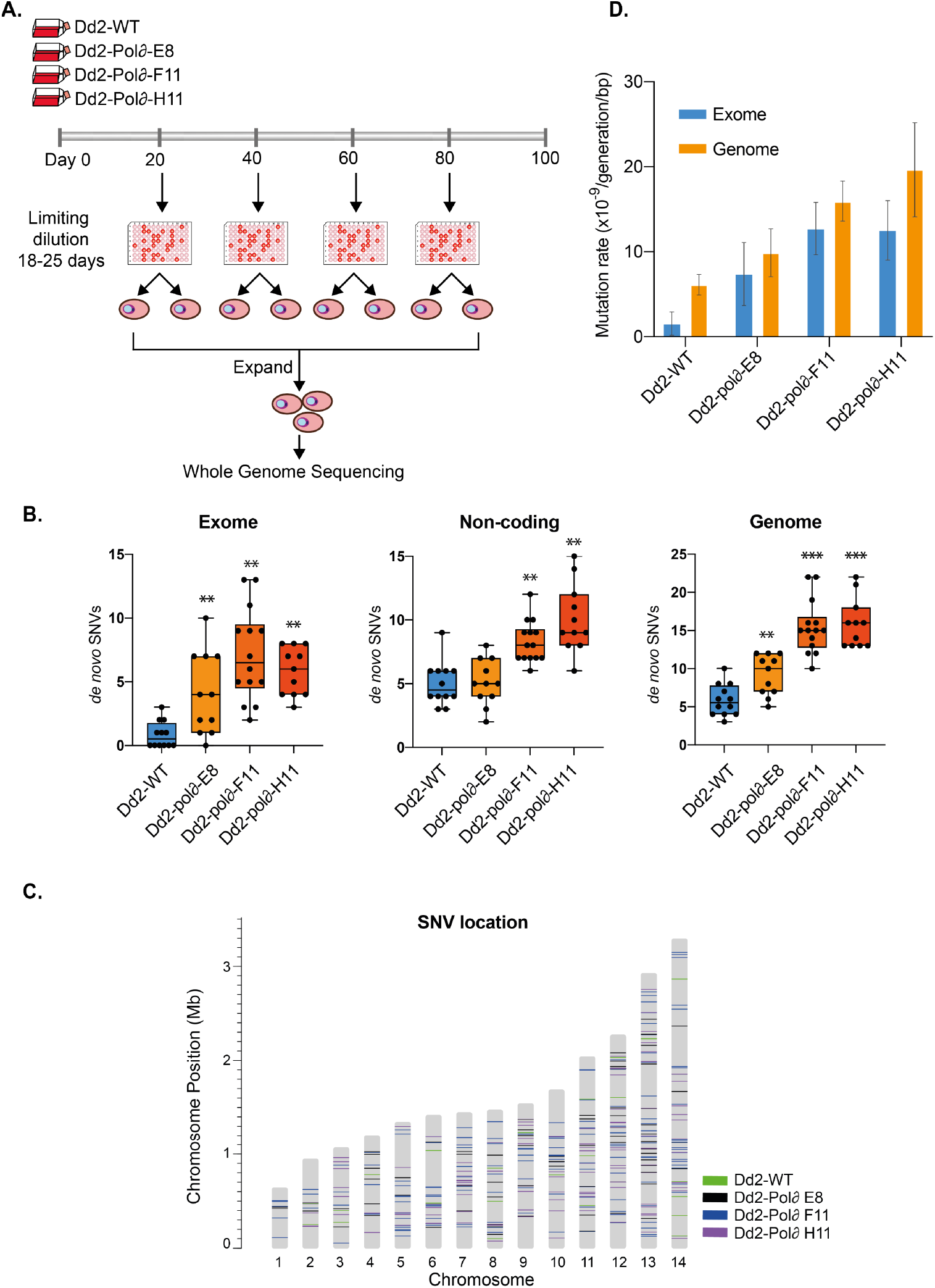
Elevated mutation rate of the DNA polymerase *δ* mutant line. **A**) Mutation accumulation assay comparing Dd2-WT with three clones of Dd2-Pol*δ*. All lines were cultured in parallel for 100 days (∼50 generations). Parasites were sampled for clonal isolation every 20 days and subsequently harvested for genomic DNA extraction. Whole-genome sequencing was performed on samples collected on day 0, 20, 40, 60, 80 and 100. **B**) The number of unique SNVs in the exome, non-coding and core genome regions of Dd2-WT and Dd2-Pol*δ* lineswere identified by subtracting from the SNVs found on day 0. Each point represents one clone, whiskers showing min-max. Wilcoxon matched-pairs signed-rank test showed statistically significant differences for the Dd2-Pol*δ* clones relative to Dd2-WT (**p<0.01, ***p<0.001). **C**) Genomic position of SNVs, colour-code by parasite line. **D**) The mutation rates of Dd2-WT and Dd2-Pol*δ* clone E8, F11 and H11, error bars showing 95% confidence intervals (data shown in Table 1).

We next determined the mutation rates for Dd2-WT and the three Dd2-Polδ clones E8, F11 and H11 based on the number of *de novo* SNVs (**Figure 2D, Table 1 and Supplementary Table 3**). Each of the Dd2-Polδ clones showed higher mutation rates than Dd2-WT, varying from 2-3 fold in the core genome (coding and non-coding regions) to 5-8 fold in coding regions (exome). Dd2-Polδ clones F11 and H11 showed a higher mutation rate than clone E8 (**Figure 2D and Table 1**), and thus all subsequent experiments were performed with clone H11. To examine whether the modest differences in mutation rate between clones might be attributed to spontaneous mutations in DNA repair genes, we also looked at whether genes playing a role in DNA repair were mutated in the Dd2-Polδ lines during the 100-day culture period (**Supplementary Table 2**). Although SNVs within or near DNA repair genes were observed in each of the Dd2-Polδ lines, no one SNV was shared among all clones. Dd2-Polδ clone E8 possessed SNVs in the coding region of two putative DNA repair genes: a G435E change in DNA polymerase theta (PfDd2_130037000) and a N420K change in DNA repair protein RHP16 (PfDd2_120056000). Dd2-Polδ clone F11 had a P225L change in RuvB-like helicase 3 (PfDd2_130068000). Dd2-Polδ clone H11 did not have SNVs in the coding region of any DNA repair genes, however, a SNV was observed in the non-coding region in proximity to proliferating cell nuclear antigen 2 (PfDd2_120031600) (**Supplementary Table 2**).

**Table 1.** The mutation rate per haploid genome or exome per generation in *P. falciparum* Dd2-WT and Dd2-Polδ parasites cultured in the absence or presence of drug pressure. Fold change was calculated by using untreated Dd2-WT as a comparator in both the no-drug and drug-pressured conditions. Values in brackets represent 95% confidence intervals, with negative values adjusted to zero.

| Parasite lines | Number of clones | Mutation rate (Genome) | Mutation rate (Exome) | Fold-change (Genome) | Fold-change (Exome) |
| --- | --- | --- | --- | --- | --- |
| **No drug pressure** |  |  |  |  |  |
| Dd2-WT | 12 | 6.12E-9 (4.91e-9 – 7.33E-9) | 1.54e-9 (1.58E-10 – 2.92E-9) | - | - |
| Dd2-Polδ-E8 | 12 | 9.86E-9 (7.06e-9 – 1.27E-8) | 7.37e-9 (3.67E-9 – 1.11E-8) | 1.6 | 4.8 |
| Dd2-Polδ-F11 | 14 | 1.59E-8 (1.36E-8 – 1.83E-8) | 1.27E-8 (9.64E-9 – 1.58E-8) | 2.6 | 8.2 |
| Dd2-Polδ-H11 | 11 | 1.97E-8 (1.41E-8 – 2.52E-8) | 1.25E-8 (9.04E-9 – 1.6E-8) | 3.2 | 8.1 |
| **With drug pressure** |  |  |  |  |  |
| **Dd2-WT** |  |  |  |  |  |
| KAE609 | 2 | 4.42E-09 (0 – 4.19E-8) | 4.64E-09 (0 – 2.99E-8) | 0.7 | 3.0 |
| **Dd2-Polδ-H11** |  |  |  |  |  |
| KAE609 | 3 | 3.72E-8 (2.63E-8 – 4.81E-8) | 4.42E-8 (1.11E-8 – 7.72E-8) | 6.1 | 28.7 |
| MMV665794 | 6 | 2.56E-8 (2.05E-8 – 3.08E-8) | 2.99E-8 (2.25E-8 – 3.73E-8) | 4.2 | 19.4 |
| Salinopostin A | 6 | 2.34E-8 (1.86E-8 – 2.82E-8) | 2.61E-8 (2.03E-8 – 3.19E-8) | 3.8 | 16.9 |
| KM15HA | 2 | 1.58E-8 (0 – 7.19E-8) | 1.93E-8 (4.88E-9 – 3.37E-8) | 2.6 | 12.5 |

### Dd2-Polδ potentiates *in vitro* drug resistance selections

Based on the assumption that a more diverse genetic repertoire available to the Dd2-Polδ cultures would increase the efficiency of selecting for drug-resistant parasites, we performed a proof-of-concept experiment comparing Dd2-WT with Dd2-Polδ using a drug with a well-characterised mode-of-action. KAE609 (cipargamin), currently in Phase II clinical trials, targets the *P. falciparum* P-type sodium ATPase 4 gene (*Pfatp*4, PF3D7_1211900) with SNVs known to confer resistance ^22^. An *in vitro* drug resistance selection was performed with a range of parasite inocula from 2×10^6^, 2×10^7^, 2×10^8^ and 1×10^9^ cells, cultured in the presence of 2.5 nM (∼5-fold IC_50_) KAE609 in three independent flasks.

After 5 days of drug treatment no viable parasites were detected by microscopy. Recrudescence of Dd2-WT was only observed with the highest starting inoculum of 1×10^9^, with parasites observed on day 18, 21 and 30 in the three independent selection flasks. In contrast, the Dd2-Polδ line returned parasites by day 12, and with a lower starting inoculum **(Figure 3A)**. All three flasks with 2×10^8^ and 1×10^9^ parasites were positive, and one out of three flasks with 2×10^7^ parasites also showed recrudescent parasites on day 12. No parasites were detected with the starting inoculum of 2×10^6^ in either line **(Figure 3A)**.

**Figure 3.**
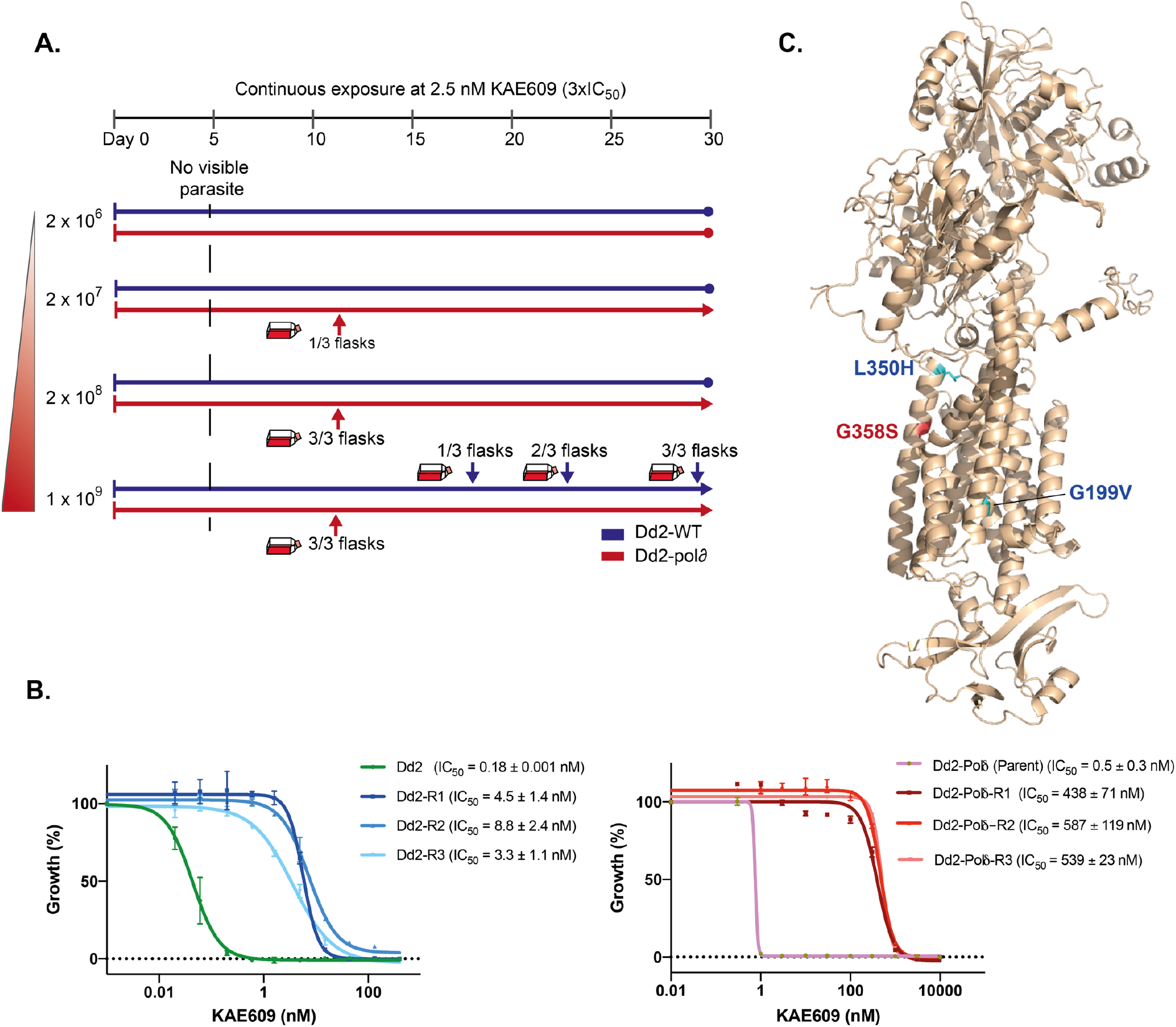
Efficient selection of resistance to KAE609 using the DNA polymerase *δ* mutant parasite. **A**) Dd2-WT (blue line) and Dd2-Pol*δ* (red line) were continuously cultured in the presence of 2.5 nM KAE609 (5×IC_50_). Parasite inocula ranged from 2×10^6^ to 1×10^9^ cells, in triplicate flasks, and parasites were detected by microscopy over the 30-day selection period. Dd2-Pol*δ* parasites were observed on day 12 with the starting inoculum of 2×10^7^, 2×10^8^ and 1×10^9^, whereas Dd2-WT parasites were detected only at the 10^9^ inoculation size, appearing in three flasks on day 18, 21 and 30, respectively. **B**) Dose-response curves of KAE609 for parental lines not exposed to drug pressure and drug-selected lines (R1-R3) for Dd2-WT (*left panel*) and Dd2-Pol*δ* (*right panel*). Shown is a representative assay (with two technical replicates, error bars showing SD), with IC_50_ ± SD values derived from three biological replicates. **C**) AlphaFold model of PfATP4 showing KAE609 resistance mutations located in or near the transmembrane domains. Blue residues originated from Dd2-WT selections, red from Dd2-Pol*δ*.

Prior to selection both the Dd2-WT and Dd2-Polδ parental lines had a similar IC_50_ of about 0.2-0.5 nM. The KAE609-selected lines from the Dd2-WT background had IC_50_ values in the range of 3 – 9 nM **(Figure 3B)**. In comparison, the drug-selected lines from the Dd2-Polδ background were appreciably more resistant with IC_50_ values around 400 – 600 nM **(Figure 3B)**, three orders of magnitude higher than their parental line.

To identify the resistance determinants driving these phenotypes, the set of selected lines was examined by whole-genome sequencing as well as direct Sanger sequencing of the *pfatp4* gene. Both approaches revealed mutations in *pfatp4* (PF3D7_1211900) in these resistant lines, with mutations at L350H and G199V in the Dd2-WT background from flask 1 and flask 3, respectively, and G358S in all lines from the Dd2-Polδ background **(Supplementary Table 4)**. All three mutations are predicted to be located within or near the PfATP4 transmembrane region ^33^ (**Figure 3C**). The mutations L350H and G358S were previously reported from an *in vitro* resistance experiment in Dd2 and 3D7 respectively with the dihydroisoquinolone SJ733, another compound targeting PfATP4 ^34^. L350H was also selected using KAE609 using a Cambodian isolate ^23^.

These results confirmed that the Dd2-Polδ line can select for drug-resistant parasites in the expected molecular target ^22, 23, 34^, with lower numbers of starting parasites (2×10^7^ vs 1×10^9^) and in a shorter period of selection than Dd2-WT (12 days vs 18-30 days).

### Dd2-Polδ successfully yields resistant parasites from an “irresistible” compound

We next challenged the Dd2-Polδ line with an “irresistible” compound. The “irresistible” class of compounds generally refers to compounds that fail to yield a drug-resistant parasite during *in vitro* selections. Identifying the mechanism of action of these compounds is of high interest due to their low propensity for resistance ^35^. MMV665794, also known as TCMDC-124162 (2-N,3-N-bis[3-(trifluoromethyl)phenyl]quinoxaline-2,3-diamine), is a quinoxaline scaffold antimalarial identified from a phenotypic high-throughput screen ^36^. Initial *in vitro* drug resistance selections were performed with this compound in wild-type 3D7 and Dd2 using different approaches, but without success **(Supplementary Table 4)**.

We treated Dd2-Polδ and Dd2-WT with 95 nM (1×IC_50_) of the quinoxaline compound intermittently. To maximise the chance of obtaining a resistant line, we used a high starting inoculum of 1×10^9^ parasites per flask, in triplicate **(Figure 4A)**. After 10-12 days of pressure, no parasites could be detected by microscopy for either line, and drug pressure was removed after day 20. Dd2-WT did not recover during the 60-day exposure period **(Figure 4A)**, consistent with previous unsuccessful selections **(Supplementary Table 4)**. In contrast, all three flasks of the Dd2-Polδ line recovered on day 21. The drug concentration was then increased to 2×IC_50_, resulting in a suppression of parasites. At day 40, cultures were switched to drug-free complete medium, and on day 60, parasites were again detected in all three flasks. Clonal lines isolated from the drug-selected parasites had an increased IC_50_ of about 2 – 2.5 fold compared with the parental line not exposed to drug pressure **(Figure 4B)**. The parasites from two flasks proliferated normally when re-exposed to constant drug pressure at 2×IC_50_, but parasites in the third flask did not survive.

**Figure 4.**
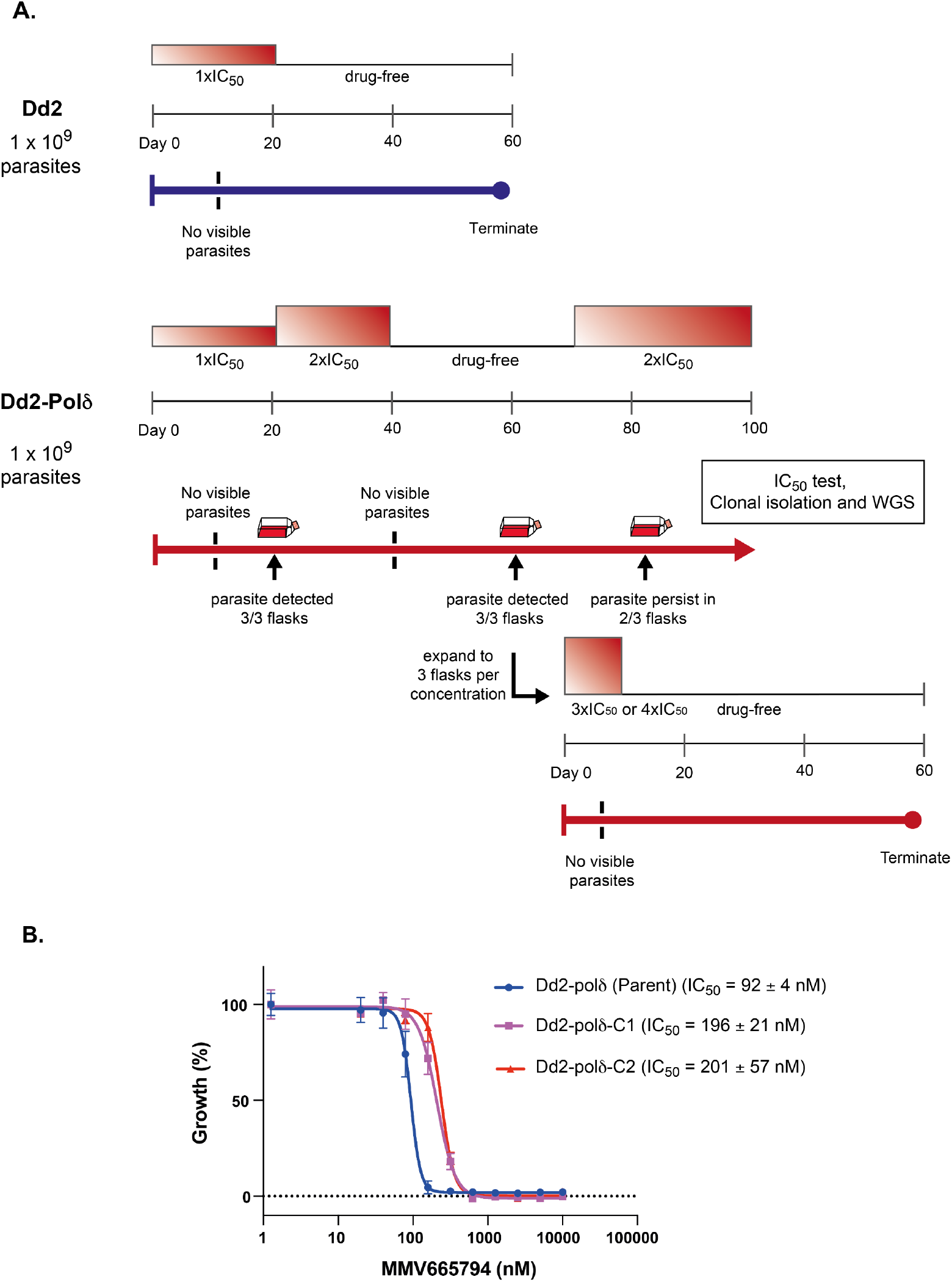
Evolution of resistance to an “irresistible” compound. **A**) Selection scheme showing inability to evolve resistance to MMV665794 with Dd2-WT, but successful isolation of resistance with Dd2-Pol*δ. Upper panel:* Dd2-WT exposed to 1×IC_50_ (95 nM) for 60 days. *Lower panel:* Dd2-Pol*δ* selection pressure was ramped to 2×IC_50_ (190 nM), with recrudescence observed after 60 days in 3 independent flasks, but only 2 of 3 flasks could stably grow under drug pressure. Recrudescent parasites were further challenged with 3× and 4×IC_50_ but failed to survive. **B**) Dose-response curves of MMV665794 for parental lines and the two resistant lines (C1-2) that were able to survive under 2×IC_50_ pressure. Shown is a representative assay (two technical replicates, error bars showing SD), with IC_50_ ± SD values derived from three biological replicates.

To investigate whether higher-level resistance could be obtained, the parasite cultures of two recrudescent flasks were each split into two more flasks that were further pressured at either 3×IC_50_ or 4×IC_50_. Parasites died after 4 days in both treatments and were subsequently cultured in drug-free complete medium (**Figure 4A**). However, no parasites recovered after 60 days, indicating that only low-level resistance could be obtained against MMV665794.

### Dd2-Polδ under drug pressure has an elevated mutation rate

The augmented ability of the Dd2-Polδ line to generate resistance to both KAE609 and MMV665794 was consistent with an increase in genetic diversity available for selection. We determined the mutation rate of Dd2-Polδ under drug pressure, comparing parasites selected with KAE609, MMV665794, and two additional resistance selections with the unrelated irresistible compounds Salinopostin A and KM15HA ^37, 38^. *De novo* SNVs of the drug-selected lines were determined in comparison with the parental lines used in the corresponding batch of selection experiments **(Figure 5A and Supplementary Figure 2B)**. The change of mutation rate in these lines was compared with non-pressured Dd2-WT, reflecting the combined factors of a defective proof-reading DNA polymerase δ and drug pressure. Dd2-Polδ under drug pressure displayed an increased mutation rate in coding regions of 13–28 fold, and ∼3–6 fold in the genome relative to non-drug pressured wild-type Dd2 **(Figure 5B and Table 1)**. When compared with non-drug treated Dd2-Polδ, these changes translate to an increase of ∼1.5–3.5 fold in coding regions and essentially unchanged (∼0.8–1.9 fold) across the genome. The Ts:Tv ratio of Dd2-Polδ under drug pressure was varied and did not show a discernible trend **(Supplementary Figure 5)**.

**Figure 5.**
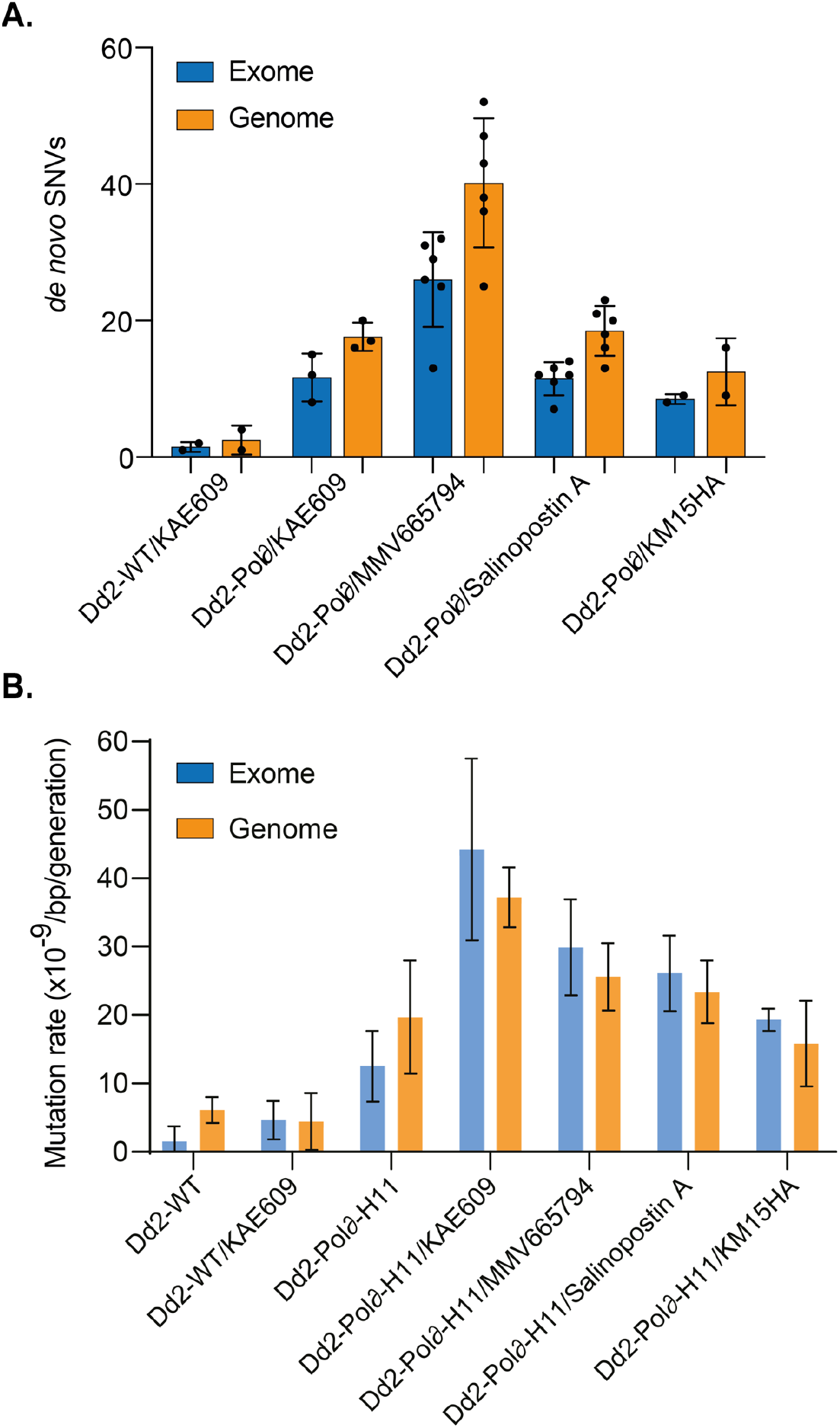
Increased number of SNVs and mutation rate of Dd2-Pol*δ* under drug pressure. **A**) The number of SNVs in Dd2-Pol*δ* selected with different antimalarial compounds. These selections, except for KAE609, failed to yield resistant parasites with Dd2-WT. Note the higher number of SNVs in KAE609-selections with Dd2-Pol*δ* compared with Dd2-WT (see Supplementary Table 6). Each dot represents a sequenced clone. The blue and orange bars represent the exome and core genome, respectively, with mean ±SD shown. **B**). The mutation rates of Dd2-WT and Dd2-Pol*δ* under drug pressure, error bars showing SD (see Table 1 for 95% confidence intervals). Data from non-drug treated Dd2-WT and Dd2-Pol*δ* clone H11 (see Figure 2D) are included for comparison.

The mutation rate of Dd2-WT under drug pressure also increased ∼3 fold in coding regions relative to non-drug pressured Dd2-WT. However this was relatively unchanged across the whole genome (**Table 1**), consistent with previous reports ^18^.

Collectively, our data indicate that the Dd2-Polδ line has an increased mutation rate that provides enhanced potential of selecting drug-resistant parasites, even with previously irresistible compounds, while being sufficiently low to maintain genome integrity and parasite robustness.

### Quinoxaline-resistant lines possess mutations in a gene of unknown function

To identify the causal resistance mutations in the MMV665794-selected lines **(Figure 4)**, we performed whole-genome sequencing on six clones isolated from two independent selections. The only mutated gene in common between all quinoxaline-selected lines was PF3D7_1359900 (PfDd2_130065800), encoding a conserved *Plasmodium* membrane protein of unknown function. The protein of 2126 amino acids encodes four predicted transmembrane domains **(Figure 6A)**. Each of the 6 clones contained one of two distinct SNVs, either G1612V or D1863Y, (equivalent to G1616V and D1864Y in Dd2, respectively) **(Figure 6A and Supplementary Table 6)**. No new copy number variants were detected in drug-selected clones **(Supplementary Table 7)**.

**Figure 6.**
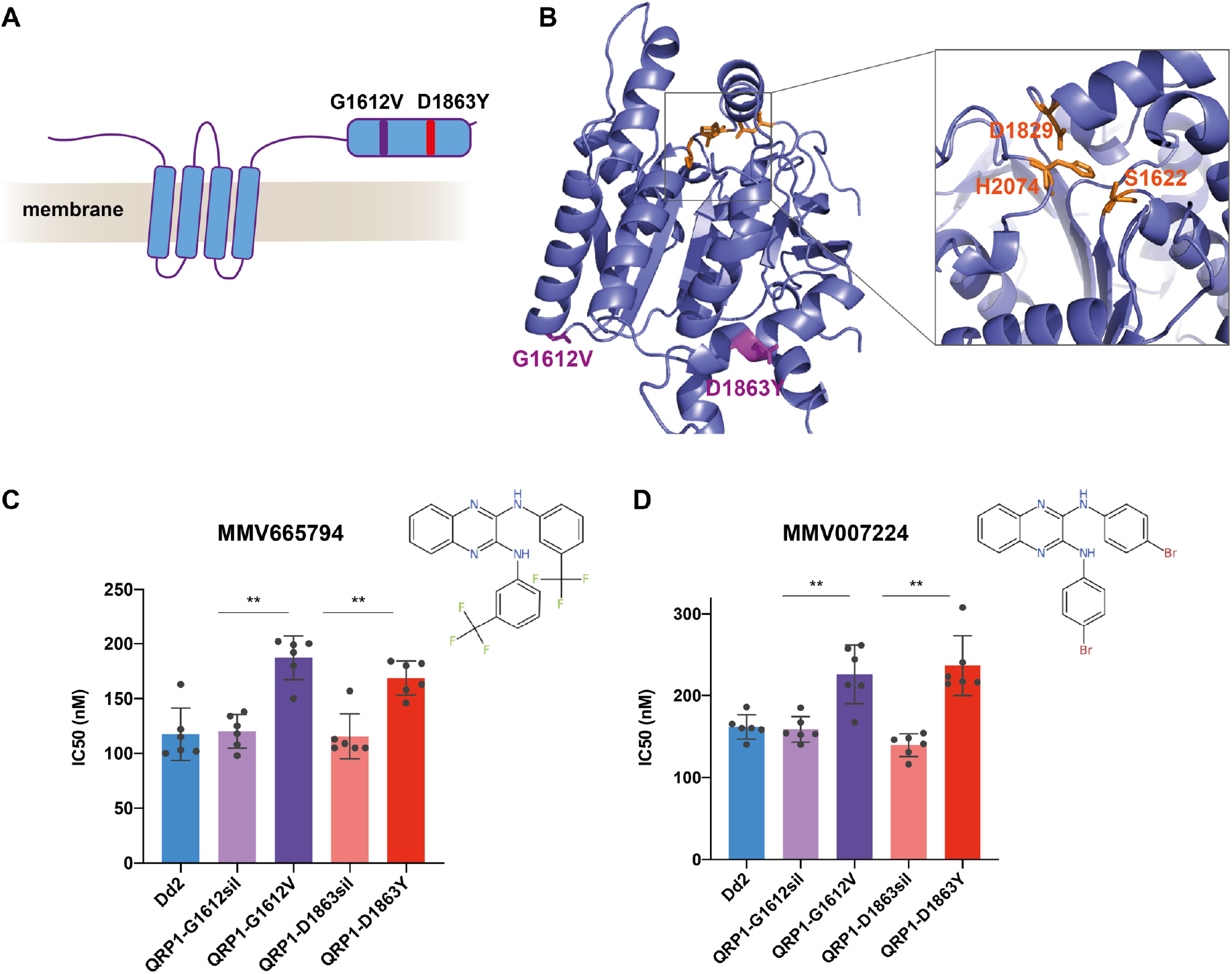
QRP1 confers resistance to quinoxaline compounds. **A**) PfQRP1 (PF3D7_1359900) encodes a 250 kDa protein with four predicted transmembrane domains. The two mutations G1612V and D1863Y found in two independent selections with MMV665794 are located near the C-terminus in a putative hydrolase domain. **B**) Model of the C-terminal 656 residues (1471-2126) of PfQRP1 showing the putative α/β hydrolase domain and catalytic triad of Ser-Asp-His (yellow), and the G1612V and D1863Y resistance mutations (purple). **C, D**) IC_50_ values of CRISPR-Cas9 edited QRP1. Dd2 lines encoding the equivalent G1612V and D1863Y mutations showed a significantly reduced susceptibility against MMV665794 and were cross-resistant to MMV007224, a structurally related molecule sharing the quinoxaline scaffold, in comparison with Dd2-WT and silent edited controls. Each dot represents a biological replicate, with mean±SD shown (**p<0.01).

To gain some insight into the potential function of PF3D7_1359900, which only has evident orthologs within the Apicomplexa (**Supplementary Figure 6**), we examined a structural model of the region containing the resistance mutations using trRosetta and AlphaFold ^39, 40^. This region is located towards the C-terminus of the protein, downstream of the 4 predicted transmembrane segments (**Figure 6A**). Protein structure comparison using the DALI server indicated potential structural homology with esterases/lipases, with a putative Ser-Asp-His catalytic triad located in close proximity on the protein model and highly conserved across orthologs (**Figure 6B and Supplementary Figure 6**).

To validate the drug-selected mutations in PF3D7_1359900, we generated CRISPR-Cas9 edited lines by introducing the Dd2 equivalent of either the G1612V or D1863Y mutation. In addition, control parasite lines were generated that were only modified with the corresponding silent mutations (G1612sil and D1863sil) and the gRNA shield mutations common to all edited lines. Both mutant lines, but not the silent controls, displayed the same modest shift in IC_50_ to MMV665794 observed in the drug selected parasites **(Figure 6B)**.

To examine whether mutations in PF3D7_1359900 had arisen in the context of other *in vitro* evolution experiments, we examined the database of SNVs identified by the Malaria Drug Accelerator Consortium in 262 *P. falciparum* lines selected with 37 compounds and identified a single clone with a frameshift mutation at residue D100 of PF3D7_1359900 ^16^. Notably, the clone had been pressured with MMV007224, a compound with a similar quinoxaline scaffold to MMV665794 **(Figure 6D)**. The presence of a frameshift mutation near the start of the protein, plus mutagenesis in the *piggyBac* whole-genome screen (Zhang et al., 2018) indicates this gene is non-essential during the asexual blood stage.

We tested whether the CRISPR-edited parasites bearing the MMV665794-resistance mutations could confer cross-resistance to MMV007224. Both the G1612V and D1863Y mutant lines showed a similar low-level resistance to MMV007224 as observed with MMV665794 **(Figure 6D)**. These findings suggest that the protein encoded by PF3D7_1359900, which we have designated as quinoxaline-resistance protein 1 (PfQRP1), may confer general resistance to quinoxaline-like compounds.

To explore whether PfQRP1-mediated resistance was specific for the quinoxaline scaffold or more broadly targets other compounds, we tested the QRP1-mutant lines against MMV665852, a compound belonging to a difference chemical class to MMV665794 and MMV007224 but that shares common pharmacophoric features of two H-bond donors linked to an aromatic ring (**Supplementary Figure 7A, B**). Although both mutant lines displayed a mildly elevated IC_50_ relative to controls, these differences were not significant. In addition, we examined a hydrolase-susceptible compound, MMV011438, which is activated by the PfPARE esterase ^41^, and GNF179 ^42^, an antimalarial expected to have an unrelated mode of action to the quinoxaline compounds. Overall, the QRP1 mutant lines did not show any significant differences for either of these compounds (**Supplementary Figure 7B**). Collectively, our data suggest that PfQP1 is a non-essential putative hydrolase that confers resistance to quinoxaline-based compounds.

## DISCUSSION

We propose that the Dd2-Polδ mutator parasite is a powerful new tool to identify targets and resistance mechanisms of antimalarials. The defective proof-reading resulting from the engineered modification to DNA polymerase δ results in an increased rate of spontaneous mutation. By expanding the genetic sequence space in cultured parasites, we reveal an enhanced capability to yield drug-tolerant parasites under *in vitro* evolution of drug resistance regimes. We observed that for selections with a drug with a known mode-of-action, KAE609 ^_22_^, we obtained resistant parasites with 10-100 fold lower inoculum and in a shorter selection window using the Dd2-Polδ line. In the case of an irresistible compound, the quinoxaline MMV665794, the Dd2-Polδ line yielded modestly resistant parasites where previously selections had failed. One potential consideration arising from the elevated mutation rate is the presence of more unrelated genetic mutations occurring during drug resistance selection. Sequencing multiple clones from more than one independent selection, coupled with genome editing validation, will therefore be important for pinpointing causal mutations.

Using a mutation accumulation assay combined with whole-genome sequencing allowed us to determine a mutation rate based on whole-genome data, not dependent on representative reporter loci ^43^. We followed wild-type and Dd2-Polδ parasites over 100 days to derive a mutation rate. For Dd2-Polδ parasites, this rate was approximately 3-fold higher than Dd2-WT across the genome, and up to 8-fold when comparing changes in the exome. Thus Dd2-Polδ requires a smaller number of parasites for a mutation to occur in its haploid genome than Dd2-WT (5.08E7 vs 1.63E8 parasites; **Supplementary Table 3**). In the presence of antimalarial compounds, the mutation rate across the genome was only modestly increased when compared with non-drug treated lines, consistent with a previous study that found a less than 3-fold mutation rate increase under atovaquone selection ^18^. However, when we consider the mutation rate in the exome after drug pressure, Dd2-Polδ was up to 28-fold higher when compared with non-drug treated wild-type parasites, and approximately 9.5-fold higher compared with drug-treated Dd2-WT (**Table 1**). This seeming increase within the exome may reflect the positive selection of functional mutations that impact the ability to survive drug pressure or to maintain fitness by supporting primary resistance mutations.

The mild mutator phenotype of Dd2-Polδ may be advantageous in two respects, by not creating too many mutations under selection to allow identification of the likely causal mutations, and by maintaining fitness despite the potential generation of detrimental mutations. In comparison with Dd2-WT, the Dd2-Polδ line showed only a minor loss of fitness, and we did not observe reversion of the engineered D308A/E310A mutations in DNA polymerase after long-term culture. In contrast, the equivalent DNA polymerase δ mutant in *P. berghei* showed a significant reduction in fitness, and the presence of an antimutator mutation in DNA polymerase δ was observed (Honma et al., 2014; Honma et al., 2016). The higher mutation rate of the *P. berghei* DNA polymerase δ mutant, approximately 90-fold over wild-type, and potentially the more stringent growth conditions *in vivo* may explain the greater impact on parasite fitness in the rodent malaria parasite. These two mutations were also not found in clinical isolates existing in the Pf6K database ^44^.

Antimutator mutations in DNA polymerases act to increase fidelity and can themselves have inherent fitness costs (Herr et al., 2011). We did not observe antimutator mutations in DNA polymerase δ in any of the sequenced *P. falciparum* Dd2-Polδ clones, perhaps a reflection of the limited selective pressure imposed by the moderate elevation in mutation rate. Nonetheless, all three Dd2-Polδ clones possessed SNVs in or near genes that play roles in DNA replication and DNA repair, although whether these mutations confer functional effects is unknown. The non-coding SNV close to proliferating cell nuclear antigen 2 (PCNA2) in clone H11 (**Supplementary Table 2**) may potentially modulate gene expression of *pcna2*, one of two PCNA proteins in *P. falciparum*, to facilitate high processivity of DNA polymerase δ ^45, 46, 47^. In addition, Dd2-Polδ clone E8, which displayed a lower mutation rate than the other two clones (F11 and H11), had a non-synonymous SNV (G435E) in the putative DNA polymerase theta (PF3D7_1331100). Gly435, equivalent to Gly226 in human DNA polymerase θ, lies in the region of the DEAD/DEAH box helicase. DNA polymerase θ possesses a low fidelity DNA polymerase and helicase activity, and plays a role in DNA repair such as double-strand break repair through canonical non-homologous end joining, microhomology-mediated end joining and homologous recombination. Polymerase θ has an impact on genome stability and repairing breaks formed by G4 quadruplex structures ^47, 48^.

The ability of the Dd2-Polδ line to elicit resistant parasites was evaluated using two compounds, KAE609 (cipagarmin) and MMV665794. KAE609, currently in Phase II clinical trials, targets the P-type ATPase PfATP4 that is responsible for transport of Na^+^ across the parasite plasma membrane (Rottmann et al., 2010; Spillman et al., 2013). All three mutations obtained from our selections were in the predicted transmembrane region of PfATP4, consistent with most previously observed mutations (Rottmann 2010; Jimenez-Diaz et al., 2014; Viadya et al., 2015; Lee and Fidock, 2016). Notably, selections with the Dd2-Polδ line yielded a G358S mutant that was recently observed in the majority of treatment failures during a Phase II trial of KAE609 (Schmitt et al., 2021), indicating that *in vitro* evolution with this line can yield outcomes with *in vivo* relevance. This high-level resistance mutation was also observed previously in selections with a dihydroisoquinolone compound (+)-SJ733 (Jimenez-Diaz et al., 2014), as well as in parallel KAE609 selections with our Dd2-Polδ line by another group (Qiu et al., 2022). Qui et al. demonstrated that the G358S mutation protects the Na^+^-ATPase activity of PfATP4 from inhibition by KAE609, but at the cost of lowering the affinity of the protein to Na^+ 49^.

In addition to obtaining more facile resistance with lower parasite numbers, the main potential of the Dd2-Polδ line is in accessing new sequence space for previously irresistible compounds. We challenged the Dd2-Polδ line with MMV665794, an irresistible compound with a quinoxaline chemical scaffold and a flutamide-like functional group ^16, 50^. Our selections with Dd2-Polδ yielded a mildly resistant parasite after approximately 60 days, whereas selections with wild-type 3D7 or Dd2 lines failed (**Supplementary Table S4**) ^16^.

Whole-genome sequencing of MMV665794-resistant clones revealed mutations in PF3D7_1359900 in all six clones selected from two independent selection flasks. The gene is predicted to be non-essential based on a single *piggyBac* insertion site approximately 0.8 kb into the 7 kb gene (Zhang et al., 2018). CRISPR-Cas9 editing of the G1612V and D1863Y mutations confirmed their role in the resistance phenotype. Furthermore, these parasites displayed cross-resistance to a structurally related quinoxaline compound, MMV007224, but not compounds from different chemical classes. The protein encoded by PF3D7_1359900, which we have termed quinoxaline resistance protein 1 (PfQRP1), is predicted to contain 4 transmembrane domains towards the N-terminus, and a putative hydrolase domain towards the C-terminus, with the resistance mutations located within the hydrolase domain region. The non-essential nature of PfQRP1 suggests this is not the target of the quinoxaline compounds but a resistance mechanism. Whether this involves direct action on the compound, in a manner akin to the PfPARE esterase (Istvan et al, 2017) or an indirect effect is not known. Nonetheless, the level of resistance elicited is modest with only a two-fold loss of potency. Thus, the difficulty in obtaining resistance to MMV665794 and a related compound, together with the limited shift in potency, suggest that compounds of this chemical class may be promising antimalarial candidates for further exploration.

Evolution of resistance *in vitro* coupled with whole-genome analysis has proven to be a highly successful technique for understanding the mechanism of action of novel compounds as well as identifying markers for drug resistance in the field ^13, 51^. One limitation of this approach has been the relative difficulty of eliciting resistance to some chemical classes. By increasing the genetic complexity of *in vitro* cultures, the Dd2-Polδ parasite line developed here has the potential to reduce the parasite inoculum, accelerate the selection time, and enable exploration of previously irresistible compounds.

## METHODS

### Genome editing using CRISPR-Cas9

*P. falciparum* Dd2 strain was employed for all genetic manipulations using CRISPR-Cas9. To generate the “mutator” line, the conserved catalytic amino acid residues D308 and E310 of DNA polymerase δ catalytic subunit (PF3D7_1017000) were mutated to alanine. Two different single guide RNAs (sgRNA) and a donor repair template harbouring the double D308A/E310A mutations with additional shield mutations to prevent sgRNA-Cas9 complex binding were cloned sequentially into a single plasmid that contains *Sp*Cas9 and the *hdhfr* selectable marker as described **(see Figure 1A)**^52^. To introduce the putative resistance mutations G1612V and D1863Y into PF3D7_1359900, two sgRNAs and a donor repair template for each mutation were constructed as above. Guide RNAs were synthesised as oligo primers (IDT). Donor repair templates and control donor templates encoding only silent mutations at the targeted sites were synthesised by GeneArt (Thermo Fisher Scientific). The 5’ and 3’ of donor DNA templates were flanked by an additional 20-21 bp sequence with homology to the destination pDC2-Cas9-gRNA plasmid for insertion at the *Aat*II and *Eco*RI sites using NEBuilder HiFi DNA Assembly ^52^. The plasmid constructs were verified by Sanger sequencing. The sgRNAs and Sanger sequencing primers are shown in **Supplementary Table 1**.

### Parasite cultivation and transfection

Parasites were cultured in RPMI 1640 (Gibco) complete medium consisting of 0.5% Albumax II (Gibco), 25 mM HEPES (culture grade), 1x GlutaMAX (Gibco), 25 µg/mL gentamicin (Gibco), and supplied with O^+^ human red blood cells (RBCs) obtained from anonymous healthy donors from National Health Services Blood and Transplant (NHSBT) or Red Cross (Madrid, Spain). The use of RBCs was performed in accordance with relevant guidelines and regulations, with approval from the NHS Cambridgeshire Research Ethics Committee and the Wellcome Sanger Institute Human Materials and Data Management Committee for the experiments performed in the UK, and sourced ethically and their research use was in accord with the terms of the informed consents under an IRB/EC approved protocol for experiments done in Spain. Parasites were routinely maintained at 0.5% – 3% parasitaemia with 3% hematocrit and were cultured under malaria gas (1% O_2_, 3% CO_2_ and 96% N_2_). A synchronous ring stage was obtained by 5% sorbitol treatment in the cycle prior to electroporation. In the next cycle, the ring stage (0–16 hours) at 10% parasitaemia was harvested for electroporation. The pDC2-Cas9-gRNA-donor plasmid was transfected into *P. falciparum* Dd2 strain using a Gene Pulser Xcell (BioRad). Fifty micrograms of plasmid was mixed with 150 µl of packed parasitised-infected red blood cells and complete cytomix (120 mM KCl, 0.2 mM CaCl2, 2 mM EGTA, 10 mM MgCl2, 25 mM HEPES, 5 mM K2HPO4, 5 mM KH2PO4; pH 7.6) to make a total volume of 420 µl ^52^. The transfectants were selected in complete medium containing 5 nM WR99210 (Jacobus Pharmaceuticals) for 8 days. The culture was subsequently maintained in drug-free complete medium until parasites reappeared. Limiting dilution was performed to isolate clonal gene-edited parasites **(Figure 1B)**. Transfectants from bulk and clonal cultures were genotyped by allele-specific PCR and Sanger sequencing. Primers are shown in **Supplementary Table 1**.

### Mutation accumulation assay

The mutation accumulation assay was performed with Dd2-WT and three Dd2-Polδ clones. Mixed-stage parasites at 1–5% parasitaemia in 10 ml were cultured continuously for 100 days. Parasites were taken out of the continuous cultures on day 0, 20, 40, 60, 80 and 100 for clonal isolation by limiting dilution in 96-well plates. One to three parasite clones from each time point were propagated for genomic DNA extraction by DNeasy Blood & Tissue Kits (Qiagen) for whole-genome sequencing on a Hiseq X (Illumina).

### Competitive fitness assay

The assay was performed by mixing the test and control parasites at a 1:1 ratio with 1% parasitaemia each. Dd2-GFP, a Dd2 line expressing green fluorescent protein from the ER hsp70 promoter ^32^, was used as the reference parasite that competed against either Dd2-WT or Dd2-Polδ in a 6-well plate. The haematocrit of the query and the competitor cultures was determined using the Cellometer Auto 1000 (Nexcelom Bioscience). Parasitaemia was determined by staining parasites with MitoTracker Deep Red FM (Invitrogen) and counting using a CytoFlex S flow cytometer (Beckman Coulter), with counts and parasite stage confirmed by microscopic examination following Giemsa staining (VWR Chemicals). Uninfected RBCs were used as a signal background for gating on the flow cytometer. The competitive fitness was determined by measuring the total parasitaemia by MitoTracker Deep Red staining, and the proportion of GFP-positive control parasites on the flow cytometer every two days for 20 days (about 10 generations). Samples were prepared in a 96-well round-bottom plate (Costar) by taking 4 µL of culture into 200 µL phosphate buffer saline (PBS) (Gibco) containing 100 nM of Mitotracker Deep Red FM. The plate was incubated at 37°C for 15 minutes and subjected to analysis on the flow cytometer. The gates were set up for the FITC (gain 5 or 10) and APC (gain 3 or 5) channels for GFP and Mitotracker Deep Red FM signals, respectively. Two independent biological experiments with three technical replicates were performed.

### *In vitro* drug resistance selections using Dd2-Polδ

Two compounds were used for *in vitro* evolution of resistance, KAE609 (cipargamin) and MMV665794, an antimalarial compound identified in the Tres Cantos Antimalarial Set and included in the Medicines for Malaria Venture Malaria Box ^36, 50^. To determine the minimum inoculum for resistance (MIR) for KAE609, three independent flasks containing ring stage cultures of Dd2-WT and Dd2-Polδ clone H11 were tested at a range of inocula ranging from 2×10^6^, 2×10^7^, 2×10^8^, and 1×10^9^ parasites. Each flask was continuously cultured in complete medium containing 2.5 nM (∼5×IC_50_) of KAE609. This concentration was able to kill parasites to a level undetectable by microscopy of Giemsa-stained thin smears. Parasite death and recrudescence after drug treatment was monitored by Giemsa staining of thin smears, with microscopic examination every day or every second day. Selections with MMV665794 were performed with Dd2-WT and Dd2-Polδ with intermittent drug exposure in three independent flasks (illustrated in **Figure 4A**). Parasites at an initial inoculum of 1×10^9^ were continuously exposed to 95 nM of MMV665794 (1×IC_50_) for 20 days. Then, Dd2-WT was maintained in drug-free complete medium until day 60. Dd2-Polδ parasites that reappeared after selection were subsequently exposed to a two-fold increment of MMV665794 at 190 nM until day 40. Drug pressure was removed until parasites were detected, and the concentration was ramped up to 3×IC_50_ and 4×IC_50_. For all selected lines, parasite clones were isolated by limiting dilution and propagated for 18–25 days. Parasites before drug pressure and surviving parasites after drug pressure were harvested for genomic DNA extraction and whole-genome sequencing.

### Drug susceptibility assay

Drug susceptibility assays were performed in 96-well plates using synchronized ring-stage parasites prepared by using 5% sorbitol. The ring stage parasites in the next cycle were diluted to 1% parasitaemia with 2% haematocrit (final concentration in the assay plate) to perform the half-maximal inhibitory concentration assay (IC_50_). The concentration range was prepared by two-fold serial dilution of compound in complete medium. KAE609 concentrations varied from 0.2–100 nM and 0.02–10 µM depending on the parasite lines. The concentration of MMV665794, MMV007224, MMV665852, GNF179 and MMV011438 ranged from 0.02–10 µM. Parasites untreated or treated with 5 µM artesunate and RBCs only (2% haematocrit) were included in the assay plate as controls. Parasite growth was determined after 72-hour drug incubation by using 2x lysis buffer containing 2×SYBR Green I (Molecular Probes) ^53^. IC_50_ analysis was performed using GraphPad Prism 8 and statistical significance was determined by Mann-Whitney *U* tests. All assays were performed in three to six independent biological experiments with two technical replicates each.

### Whole genome sequencing

Parasite samples were lysed in 0.1% saponin, washed with 1×PBS, and genomic DNA (gDNA) was extracted using the QIAamp DNA Blood Midi Kit (Qiagen). The concentration of gDNA was quantified using the Qubit dsDNA BR assay kit and measured with a Qubit 2.0 fluorometer (Thermo Fisher Scientific) prior to sequencing. The samples were sheared to around 450-bp fragments and the library constructed using the NEBNext UltraII DNA library kit (NEB), followed by qPCR for sample pooling and normalisation for the Illumina sequencing platform. Paired-end sequencing (2×150 bp) and PCR-free whole genome sequencing was performed on a HiSeq X (Illumina) ^54^. Samples selected for resistance to Salinopostin A and KMHA15 were sequenced on an Illumina MiSeq or NextSeq 550 sequencing platform, respectively, to obtain 300 or 150 bp paired-end reads at an average of 30× depth of coverage.

### Single nucleotide variant and copy number variant calling

The genome sequences of Dd2-Polδ clones were analysed by following the GATK4 best practice workflow ^55^. Paired-end sequencing reads from each parasite clone were aligned to *P. falciparum* 3D7 (PlasmoDB-44_Pfalciparum3D7) and Dd2 reference sequence (PlasmoDB-44_PfalciparumDd2) using bwa mem (bwa/0.7.17=pl5.22.0_2). PCR duplicates were removed by GATK MarkDuplicates (picard/2.22.2--0) (**Supplementary Table 10**). Variant calling was performed by GATK HaplotypeCaller (gatk/4.1.4.1). The SNVs had to pass the filtering criteria (ReadPosRankSum ≥ -8.0, MQRankSum ≥ -12.5, QD ≥ 20.0, SOR ≥ 3.0, FS ≤ 60.0, MQ ≥ 40.0, GQ ≥ 50.0, DP ≥ 5.0). Variants that had heterozygous calls or were located outside of the core genome were excluded (The Dd2 core genome coordinates are shown in **Supplementary Table 9**). Genetic variant annotation and functional effect prediction were determined by using snpEff ^56^. Transcription start sites were mapped according to the recent refined data set ^57^

The number of *de novo* SNVs occurring during the mutation accumulation assay was identified by using the genome of Dd2-WT and Dd2-Polδ collected on Day 0 for subtraction in each parasite line. For the drug pressure condition, the *de novo* SNVs were identified by using the genome of the parental line that was not exposed to drug pressure for subtraction. The significant change of SNV numbers in Dd2-WT and Dd2-Polδ in the condition without drug was determined by Wilcoxon matched-pairs signed-rank tests (GraphPad Prism 8).

CNVs were detected by the GATK 4 workflow ^58^ adapted for *P. falciparum* as described ^59^. Briefly, read counts were collected across the genic regions of the *P. falciparum* core genome ^60^ and denoised log_2_ copy ratios were calculated against a panel of normals constructed from non-drug-selected Dd2 samples. CNVs were retained if at least 4 sequential genes showed a denoised log_2_ copy ratio greater than or equal to 0.5 (copy number increase) or less than or equal to -0.5 (copy number decrease).

### Mutation rate determination

The mutation rate (µ) of each parasite line was determined by the mean number of *de novo* single nucleotide variants (S) from all clones (C) that occurred during continuous parasite culture and that differed from the parasite line on Day 0. The duration of erythrocytic life cycles (L) and Genome size (G) were calculated as shown below ^20, 21^. A single asexual blood stage cycle for Dd2 was calculated at 44.1 hours ^20^. The sizes of the Dd2 core genome and coding region were set as 20,789,542 bp and 11,553,554 bp, respectively **(Supplementary Table 2)**. Shapiro-wilk normality test was used to examine SNV datasets for normal distribution. One sample t-test was used to examine mean samples and 95% confidence intervals **(Supplementary Table 3)**. All tests were run by R programming.

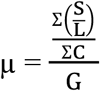

### Protein structure modelling

The protein structures of PfATP4 (PF3D7_1211900) and PfQRP1 (PF3D7_1359900) were modelled by AlphaFold ^39^ and comparisons performed using the DALI server ^61^. Structures were displayed using PYMOL molecular graphics system.

## Data availability

The data underlying this article are available within the supplementary material files. All associated sequence data are available at the NCBI Sequence Read Archive under accession code ERP110649 (BioProject: PRJEB2844). Library names DN581642P:A7, D7, E7 and DN573783H:A5-12, B5-B12, C5-C12, D5-D8, D10-D12, E5-E12, F5-F12, G6-G7, G9-G12, H4-H12.

## Acknowledgements

We would like to thank current and former members of the Lee lab for constructive feedback. We are grateful to Liz Huckle and the staff in Sanger Scientific Operations for their support with sequencing. We thank Kotanan N. for artwork. We would like to acknowledge funding from the Bill and Melinda Gates Foundation to MCSL, DAF, GSK and EAW (OPP1054480), and to DAF (INV-033538). This research was funded in whole, or in part, by Wellcome [Grant number 206194/Z/17/Z] to MCSL. For the purpose of open access, the author has applied a CC BY public copyright licence to any Author Accepted Manuscript version arising from this submission.

## Author contributions

KK and MCSL conceived the study. KK generated the Dd2-Pol∂ parasite line and performed the mutation accumulation experiments. KK, KAS, SM, VH and MGG performed the *in vitro* drug selection experiments. Drug sensitivity assays were performed by KK, KAS and MCSL. HP and ACU contributed to the generation of whole-genome sequencing data. Whole-genome sequencing data were analysed by KK, TK, TY, ML, RP, JH and SM. FJG, EAW, DAF, TC and MCSL planned the experiments and supervised the study. All authors contributed to writing the paper.

## List of Supplementary Material

### Supplementary Figures

**Supplementary Figure 1**. Alignment of DNA polymerase *δ* from different species

**Supplementary Figure 2**. *De novo* SNVs observed in the absence and presence of drug pressure

**Supplementary Figure 3**. Genomic position of *de novo* SNVs.

**Supplementary Figure 4**. Transition:transversion (Ts:Tv) ratio of base pair substitutions.

**Supplementary Figure 5**. Transition:transversion (Ts:Tv) ratio of base pair substitutions in cultures exposed to drug-pressure

**Supplementary Figure 6**. Alignment of QPR1 in different apicomplexan species

**Supplementary Figure 7**. Drug susceptibility of CRISPR-edited QRP1 lines

### Supplementary Tables

**Supplementary Table 1**. Number of *de novo* SNVs in Dd2-WT and Dd2-Polδ-MT occurring during the mutation accumulation assay.

**Supplementary Table 2**. List of SNVs in Dd2-WT and Dd2-Polδ occurring during the mutation accumulation assay identified by whole genome sequencing.

**Supplementary Table 3**. Mutation rate calculation

**Supplementary Table 4**. Summary of unsuccessful MMV665794 selections using wild type 3D7 and Dd2 strains

**Supplementary Table 5**. The number of *de novo* single nucleotide variants in Dd2-Polδ-H11 occurring in coding and non-coding regions during *in vitro* drug resistance selections.

**Supplementary Table 6**. SNVs in drug-selected lines in the Dd2-Polδ-H11 identified by whole genome sequencing.

**Supplementary Table 7**. CNV analysis: Denoised Log_2_ copy ratios for MMV665794-selected clones.

**Supplementary Table 8**. Single guide RNAs and sequencing primers for verifying CRISPR plasmid constructs and CRISPR-edited parasites.

**Supplementary Table 9**. The coordinates of the Dd2 core genome, translated from 3D7 core genome coordinates.

**Supplementary Table 10**. The unique and duplicate reads from WGS and % of genome having read depth > 10 and > 5.

