## Supplementary Figures 1-7 for "Generation of a mutator parasite to drive resistome discovery in *Plasmodium falciparum*"

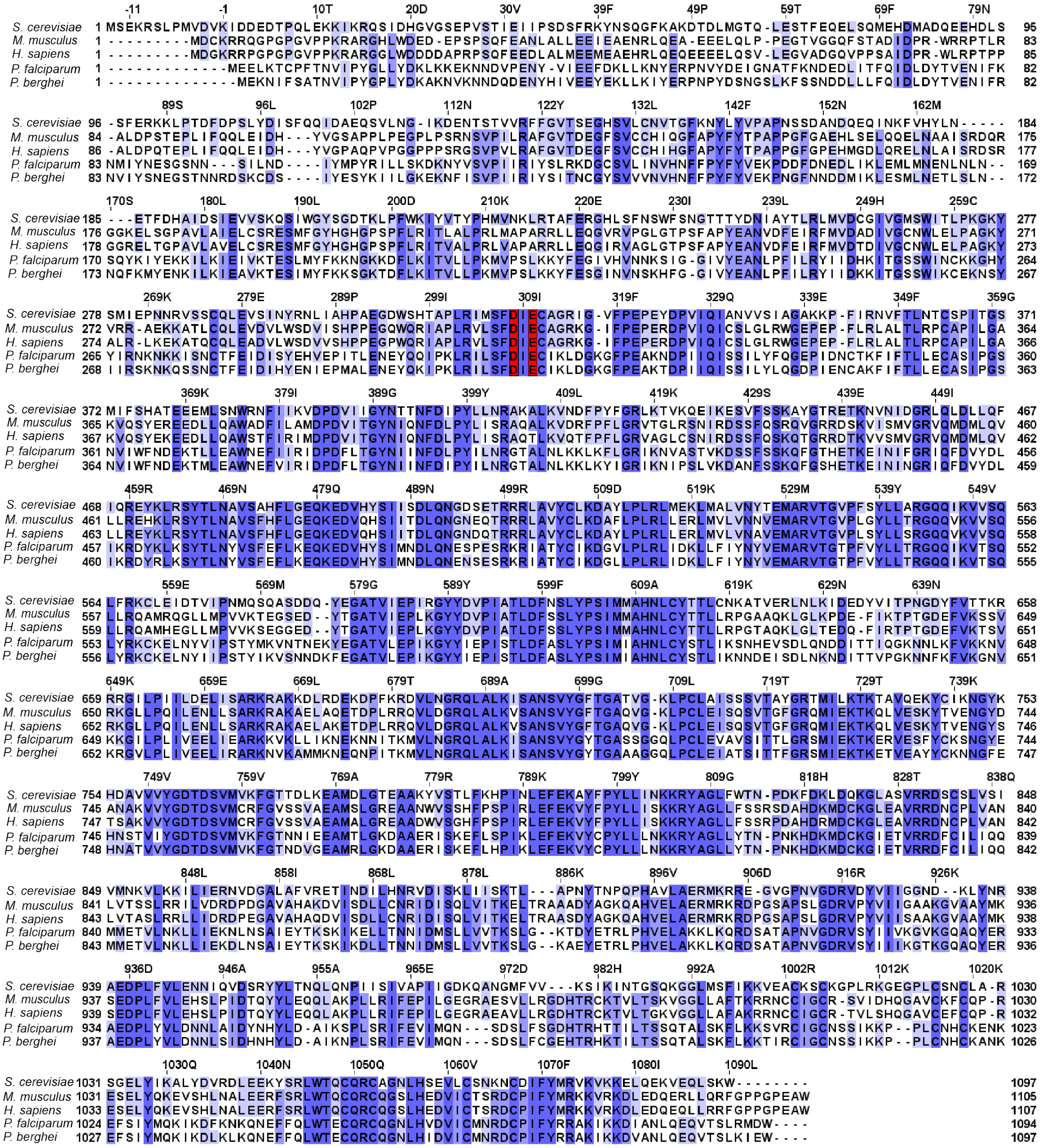


**Supplementary Figure 1. Alignment of DNA polymerase $\delta$ from different species.** The DNA polymerase $\delta$ catalytic subunit for yeast (*S. cerevisiae*), mouse (*M. musculus*), human (*H. sapiens*) and two malaria parasites (*P. falciparum* and *P. berghei*) were aligned using Clustal Omega. The two conserved catalytic residues of the 3'-5' exonuclease subunit mutated in the Dd2-Pol$\delta$ line (D308A / E310A) are highlighted in red.

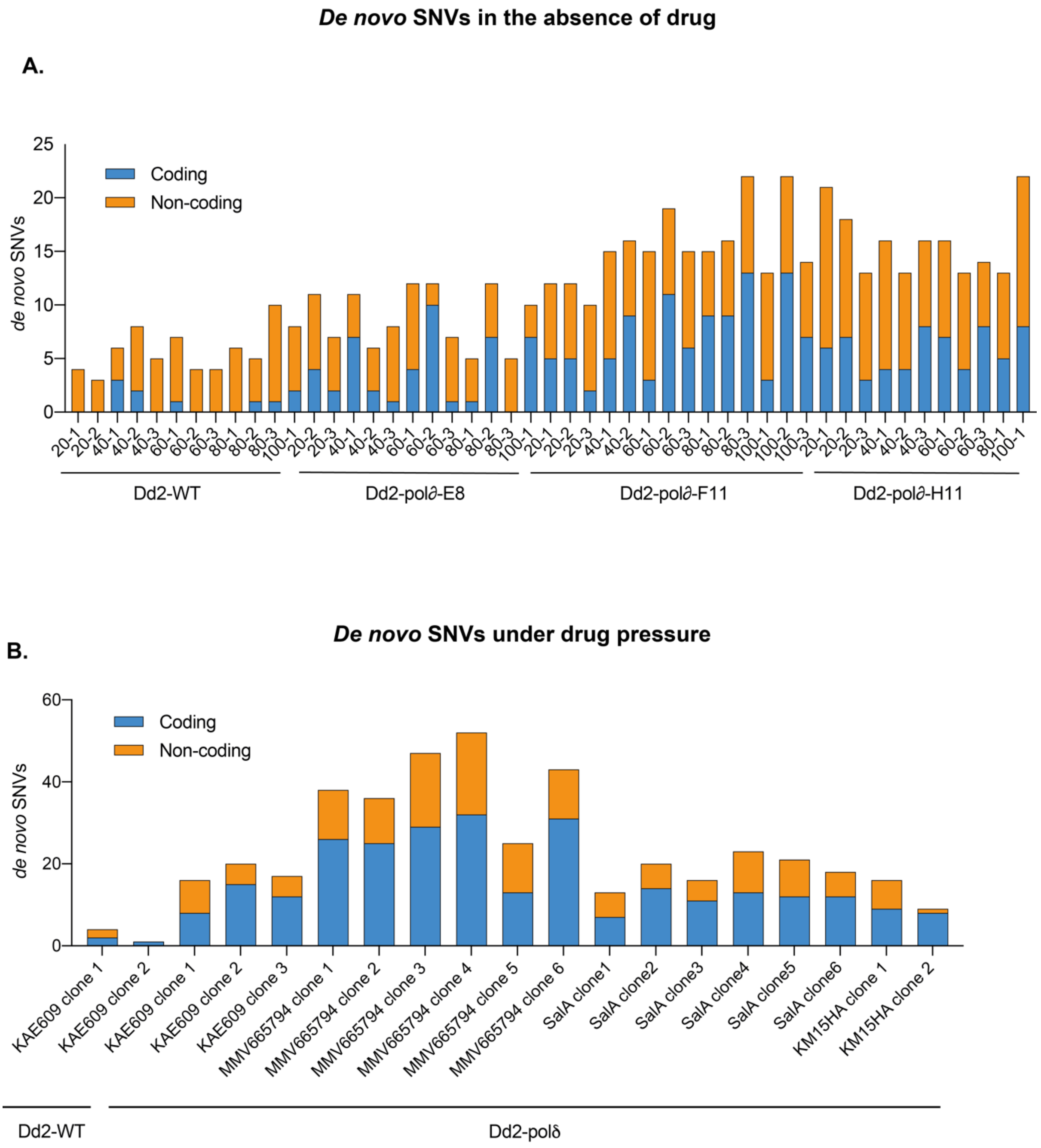


**Supplementary Figure 2. *De novo* SNVs observed in the absence and presence of drug pressure.** **A**) SNVs observed during the mutation accumulation assay performed with Dd2-WT and three different clones of Dd2-Pol$\delta$. Parasite lines were grown in continuous culture for 100 days and sampled for whole-genome sequencing (relates to Figure 2). Labels refer to the collection day and clone number (e.g. 20-1: day 20 clone 1). **B**) SNVs observed in clonal drug-evolved parasites (relates to Figure 5 and Supplementary Table 6).

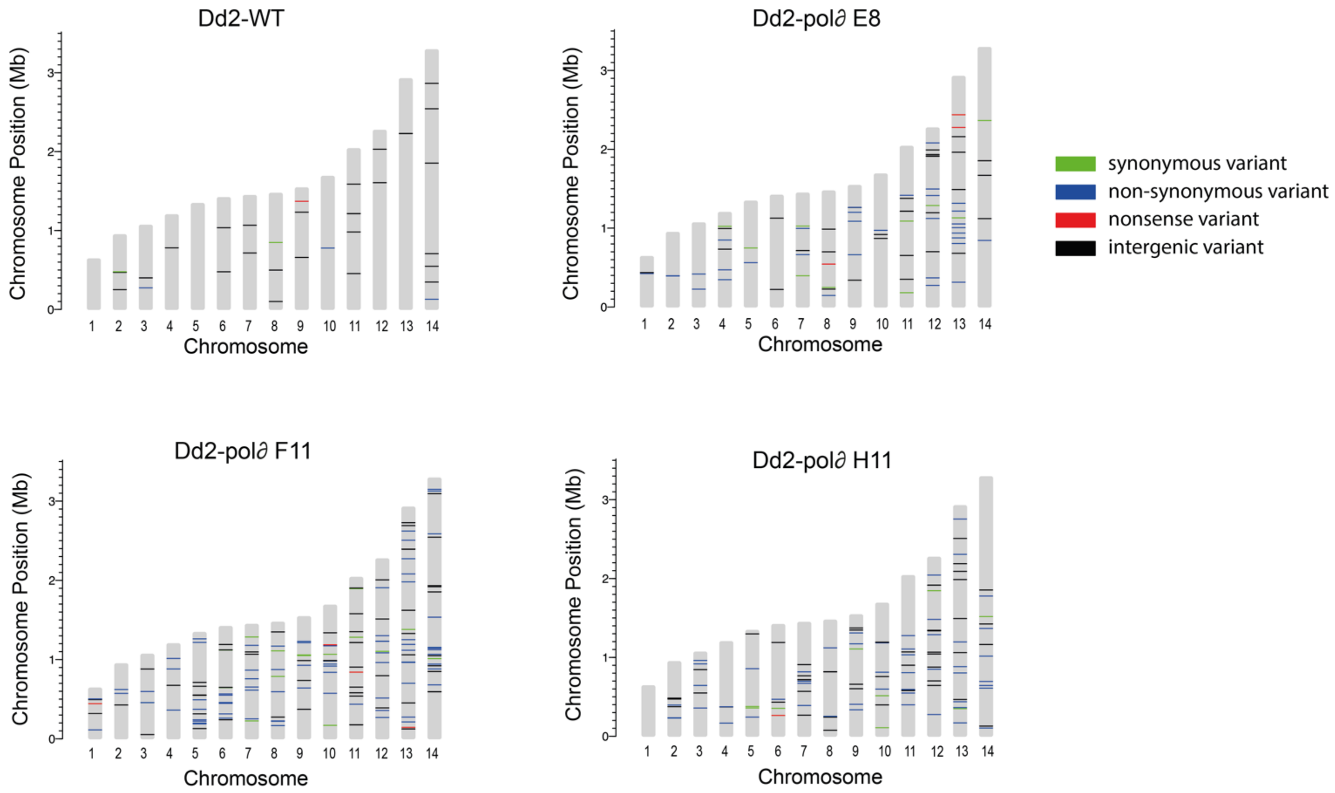


**Supplementary Figure 3. Genomic position of *de novo* SNVs.** SNVs observed in the mutation accumulation experiment (Figure 2) are displayed for Dd2-WT and the three Dd2-Pol$\delta$ clones. Dashed lines indicate their position on each chromosome, and colors indicate the mutation type.

**A.**

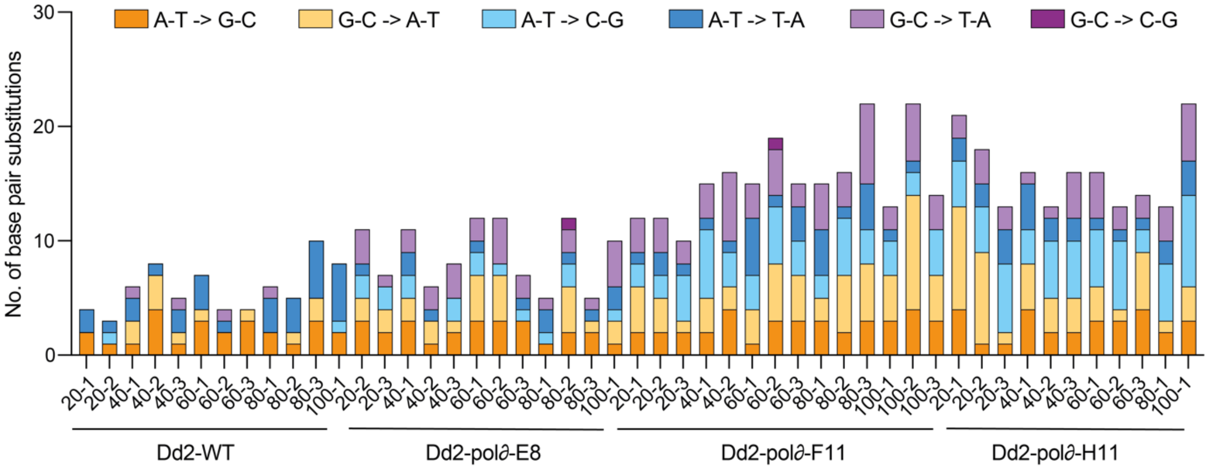


**B.**

| | | Base pair substitutions | | | | | |
|---|---|---|---|---|---|---|---|
| | | Transitions | | Transversions | | | |
| Parasite line | Ts:Tv | A:T > G:C | G:C > A:T | A:T > T:A | G:C > T:A | A:T > C:G | G:C > C:G |
| Dd2-WT | 1.04 | 2 | 0.9 | 2.3 | 0.3 | 0.2 | 0 |
| Dd2-Polδ-E8 | 0.90 | 2.2 | 2 | 1 | 2.3 | 1.3 | 0.08 |
| Dd2-Polδ-F11 | 0.72 | 2.6 | 3.9 | 1.9 | 3.6 | 3.4 | 0.07 |
| Dd2-Polδ-H11 | 0.66 | 2.6 | 3.7 | 2.1 | 2.6 | 4.8 | 0 |

**Supplementary Figure 4. Transition:transversion (Ts:Tv) ratio of base pair substitutions.** The base pair substitutions for transition (A:T → G:C and G:C → A:T) and transversion (A:T → T:A, G:C → T:A, A:T → C:G, and G:C → C:G) were examined in the Dd2-WT and Dd2-Polδ clones. **A**) Base pair substitutions in Dd2-WT and three Dd2-Pol*δ* lines in drug-free media over 100 days. Labels refer to the collection day and clone number (e.g. 20-1: day 20 clone 1). **B**) The number of base-pair substitutions were averaged for clones of each line collected during the 100-day assay. The Ts:Tv ratio for Dd2-WT was 1.04, whereas the Ts:Tv ratio for the Dd2-Polδ clones ranged from 0.66 - 0.90. This moderately decreased Ts:Tv in Dd2-Polδ indicated that base pair substitutions tend towards transversions. This was evident especially for base-pair changes from G:C → T:A and A:T → C:G that showed an increased frequency of 7-12 fold and 5-24 fold, respectively. In addition, the transitions from G:C → A:T in Dd2-Polδ increased about 2-4 fold.

**A.**

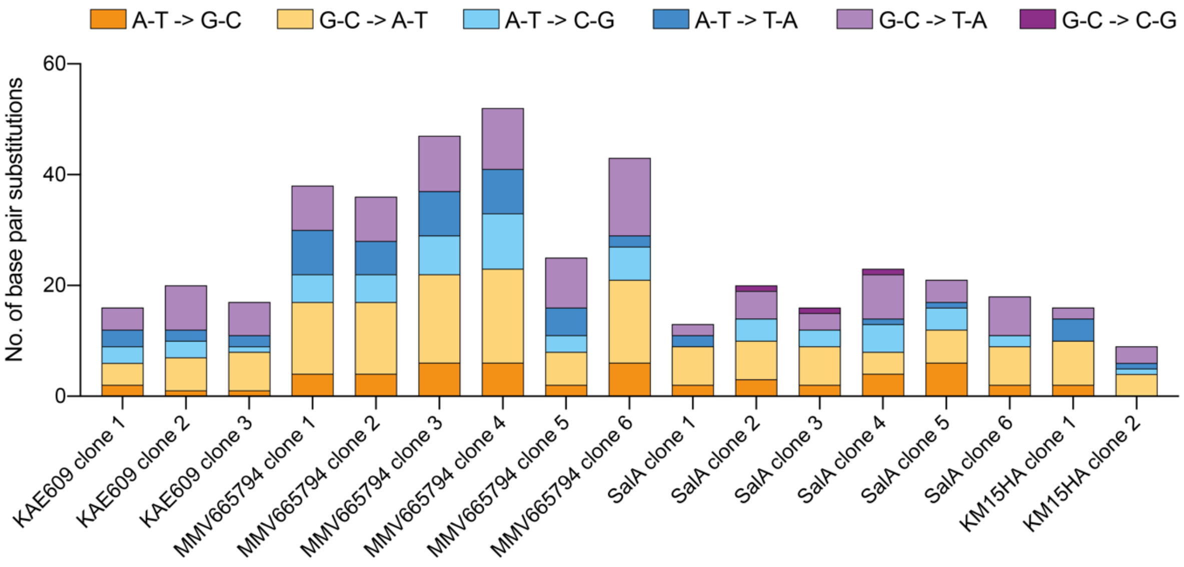


**B.**

| | | Base pair substitutions | | | | | |
|---|---|---|---|---|---|---|---|
| | | Transitions | | Transversions | | | |
| Compound | Ts:Tv | A:T > G:C | G:C > A:T | A:T > T:A | G:C > T:A | A:T > C:G | G:C > C:G |
| KAE609 | 0.68 | 1.3 | 5.7 | 2.3 | 6.0 | 2.3 | 0 |
| MMV665794 | 0.81 | 4.7 | 13.3 | 6.2 | 10.0 | 6.0 | 0 |
| Salinopostin A | 1.05 | 3.2 | 6.3 | 0.7 | 4.8 | 3.0 | 0.5 |
| KM15HA | 1.30 | 1.0 | 6.0 | 2.5 | 2.5 | 0.5 | 0 |
| Dd2-Polδ-H11 | 0.66 | 2.6 | 3.7 | 2.1 | 2.6 | 4.8 | 0 |

**Supplementary Figure 5. Transition:transversion (Ts:Tv) ratios of base pair substitutions in cultures exposed to drug-pressure. A**) Base pair substitutions in the Dd2-Polδ clone H11 after *in vitro* evolution of resistance to KAE609, MMV665794, Salinopostin A, and KM15HA. **B**) The number of base-pair substitutions were averaged for clones of each selection.

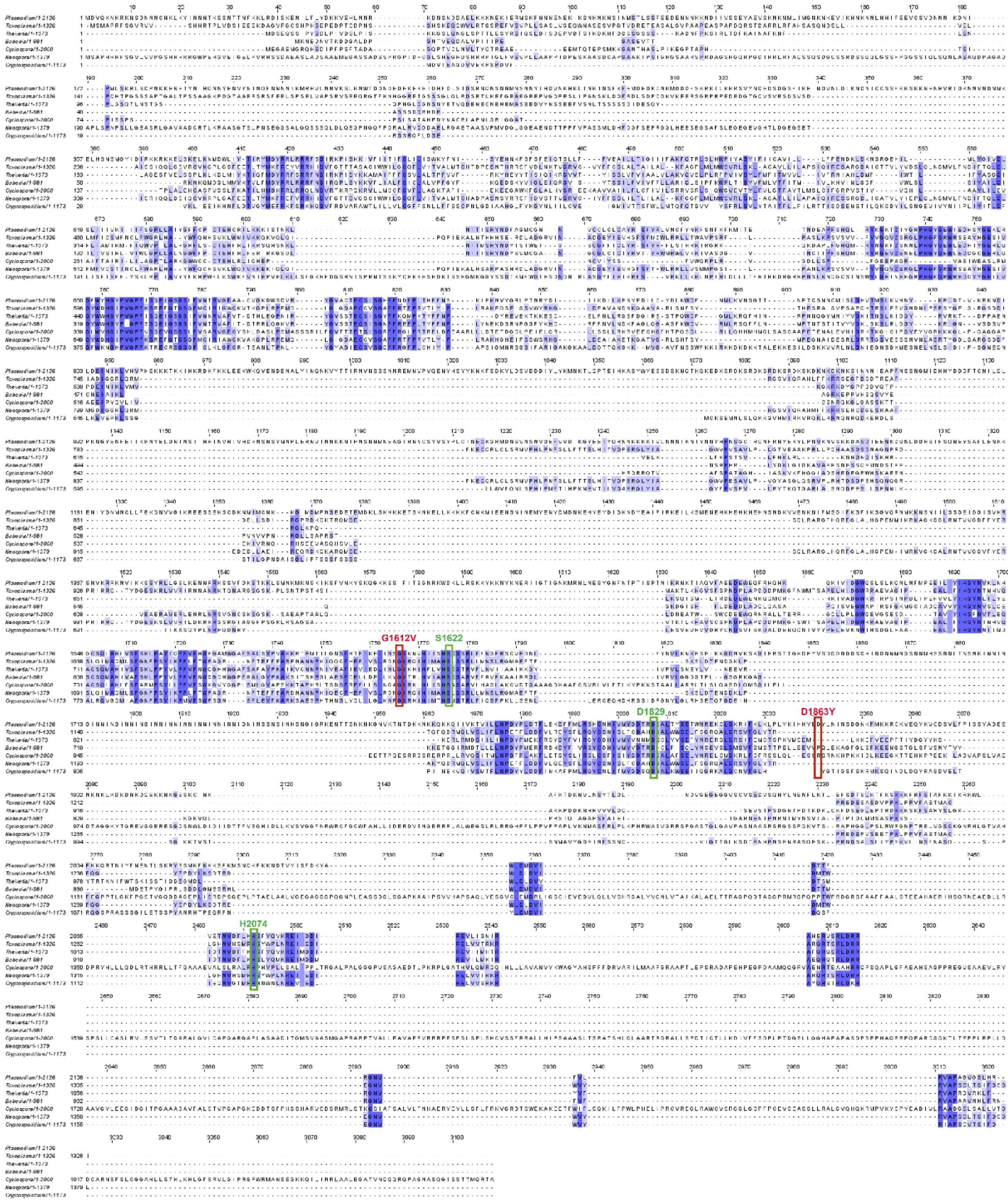


**Supplementary Figure 6. Alignment of QPR1 in different apicomplexan species.** Putative homologs of PfQRP1 (PF3D7_1359900) were identified using BLAST and aligned with Clustal Omega. Shown are *Toxoplasma gondii* (TGME49_289880), *Theileria parva* (TpMuguga_02g02080), *Babesia bovis* (BBOV_II004840), *Neospora caninum* (NCLIV_042110), *Cyclospora cayetanensis* (cyc_01400) and *Cryptosporidium parvum* (cgd3_590). Mutations involved in quinoxaline resistance are shown in red (G1612V & D1863Y), and the putative catalytic triad (Ser-His-Asp) is highlighted in green.

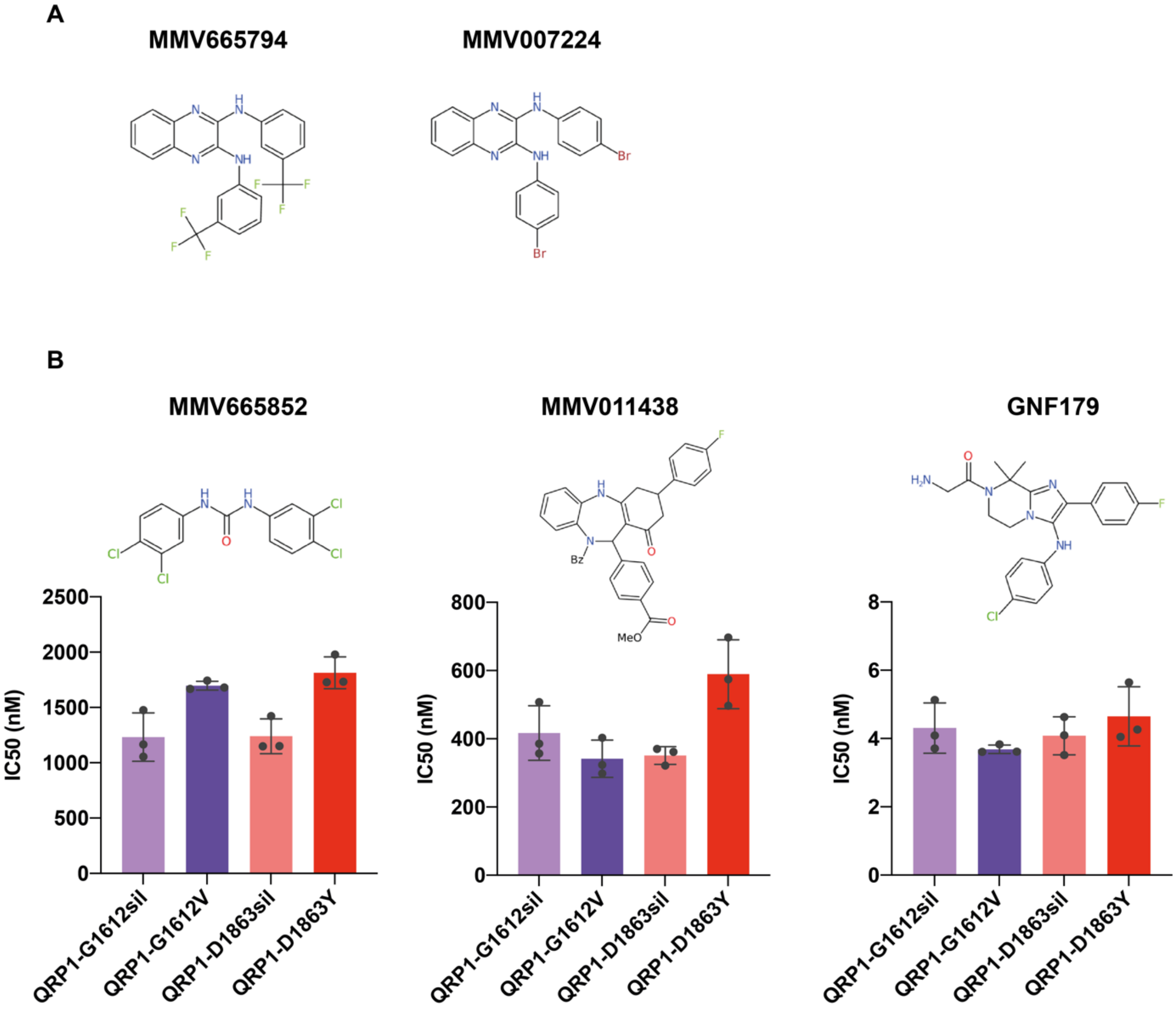


**Supplementary Figure 7. Drug susceptibility of CRISPR-edited QRP1 lines. A**) Structures of MMV665794 and MMV007224. **B**) $IC_{50}$ values of CRISPR-edited QRP1 lines. Parasite lines encoding the G1612V and D1863Y mutants or silent controls were tested against MMV665852 (Corey et al., 2016), MMV011483 (Istvan et al., 2017) and GNF179 (Meister et al., 2011). Each dot represents a biological replicate, with mean±SD shown. No significant differences were observed.
